# Distinguishing Closely Related Pancreatic Cancer Subtypes In Vivo by 13C Glucose MRI without Hyperpolarization

**DOI:** 10.1101/511543

**Authors:** Shun Kishimoto, Jeffrey R. Brender, Shingo Matsumoto, Tomohiro Seki, Nobu Oshima, Hellmut Merkle, Galen Reed, Albert P. Chen, Jan Henrik Ardenkjaer-Larsen, Jeeva Munasinghe, Keita Saito, Kazu Yamamoto, Peter L. Choyke, James Mitchell, Murali C. Krishna

## Abstract

Metabolic differences between patients and within the tumor itself can be an important determinant in cancer treatment outcome. However, methods for determining these differences non-invasively in vivo have been lacking. Using pancreatic ductal adenocarcinoma as a model, we demonstrate that tumor xenografts with a similar genetic background can be distinguished by their differing rates of metabolism, as detected by imaging of uniformly ^13^C labeled glucose tracers using a newly developed technique using tensor decomposition for noise suppression to bring the signal to a detectable level without hyperpolarization of the tracer. Using this method, cancer subtypes that appeared to exhibit similar metabolic profiles by other techniques that measured steady state metabolism can be distinguished.

---

Tumor cells are metabolically flexible and redirect metabolites away from the citric acid cycle to the less energetically efficient aerobic glycolytic pathway to generate the biomass required for growth.^*1*^ Beyond this metabolic phenotype, known as the Warburg effect, tumor growth is further aided by neo-angiogenesis for developing vasculature to support tumor growth. Compared to normal vasculature, which is well organized and structurally robust, the vascular network of tumors is chaotically organized and leaky resulting in poor delivery of oxygen and regions of both chronic and acute hypoxia.^*2*^ Thus, the tumor microenvironment is typically distinguished by high uptake of glucose, high glycolytic flux and hypoxia and acidic conditions. This characteristic metabolic and physiologic phenotype is used to develop diagnostic imaging methods and to tailor appropriate therapies.

While certain common features of the tumor microenvironment are retained in different cancers,^*3, 4*^ other features can vary considerably among patients and even within a single tumor itself.^*5, 6*^ Local concentrations of glucose,^*7, 8*^ fatty acids,^*9*^ and amino acids,^*8*^ for example, have been shown to influence the efficacy of specific types of chemotherapy and radiotherapy, which could lead to a possible change of treatment.^*10*^ Personalizing treatment in response to fluctuations of metabolites requires a reliable way of measuring the local concentrations of small molecules, which is less well-established than techniques for measuring the cellular, genomic, and proteomic environment. Molecular imaging techniques based on PET have been successful at characterizing the upregulated uptake of several probes in cancer, though such measurements are usually limited to steady state uptake only and usually cannot characterize downstream biochemical transformations.^*11*^ MALDI-MS has been employed to image metabolite concentrations of cancer tissue ex situ^*11, 12*^ but cannot be employed in vivo. Tumor metabolism in vivo can be imaged by ^1^H magnetic resonance spectroscopy, but as a steady state method it cannot distinguish the internal metabolism of the tumor from contributions from the surrounding stroma.

^13^C MRI can in principle distinguish metabolic processes using exogenous metabolic tracers. The ∼ 4 orders of magnitude lower sensitivity of ^13^C MRI is typically overcome by use of dynamic nuclear polarization, which takes advantage of the fact that the high spin polarization of a paramagnetic radical can be transferred to the ^13^C nucleus on another molecule under resonant microwave irradiation. However, this transfer happens efficiently only at temperatures near ∼1 K and the hyperpolarization is rapidly lost when the sample is brought to room temperature before injecting. Due to this limitation, these studies are normally applied to probes whose ^13^C T1 relaxation time is long enough that the enhanced polarization is not lost before the metabolic flux can be determined. Of these probes, pyruvate has proven one of the most useful as it interrogates the central switching point from glycolysis to the TCA cycle.^*13*^ By comparing the pyruvate-to-lactate conversion between tumors or between pre-and post-treatment, it has been possible to assess the glycolytic profile of tumors *in vivo* and assess metabolic changes during treatment.^*14*^ However, hyperpolarized MRI using pyruvate is unable to detect changes occurring upstream of the TCA cycle, which are common in many cancers.

An alternative approach that allows a more comprehensive analysis is to use glucose as a metabolic tracer.^*15, 16*^ Although glucose itself is difficult to hyperpolarize, new techniques allow the dynamic imaging of metabolic tracers by MRI without hyperpolarization. This imaging clearly suffers from a lack of signal but this can be compensated for by efficient noise suppression.^*17*^ To see how a targeted approach using hyperpolarized pyruvate compares to the more comprehensive approach offered by non-hyperpolarized glucose, we analyzed two closely related cell lines with a similar genetic background in the metabolic pathways. MiaPaCa-2 and Hs766t are cell lines established from pancreatic ductal adenocarcinomas (PDACs) with similar mutations in major metabolic genes.^*18*^ However, other differences in the anatomy of the xenografts and in non-metabolic pathways of the cell lines can impact the tumor microenvironment. Hs766t was derived from a metastatic site and is expected to have a different stromal boundary compared to MiaPaCa-2, which is derived from a primary heterogeneous tumor. Hs766t tumor xenografts are more strongly hypoxic than MiaPaCa-2,^*19*^ have a more poorly developed vasculature system and was expected to have a different overall physicochemical environment as a result of this anatomical difference. While MiaPaCa-2 and Hs766t have similar metabolism overall,^*10*^ it is possible to detect a difference in glucose metabolism using a newly developed technique to image glycolysis using non-hyperpolarized ^13^C glucose as a tracer.^*17*^ Imaging of local metabolite concentrations and biochemistry in this manner may provide a new method for understanding the tumor biochemical microenvironment.

## Results

### MiaPaCa-2and Hs766t PDAC Xenografts have Distinct Anatomical and Histological Characteristics

Figs 1A and C show transverse slices from the anatomical T2-weighted RARE MRI of xenografts of Hs766t and MiaPaCa-2. Both tumors are poorly differentiated and show the gross anatomy typical of Grade 3 PDACs.^*20*^ While the gross anatomy is similar, the anatomical microstructure and histology of the two tumors is distinctly different. The MiaPaCa-2 tumors appear entirely homogenous and undifferentiated, an observation that holds down to the cellular level (Fig 1B). By contrast, the homogeneity of the Hs766t tumors is broken by hypointense spots, a feature characteristic of focal necrosis (Fig 1D).^*21*^ As noted in previous reports,^*2*^ we found similar levels of CD31, a common biomarker for angiogenesis (Fig S1),^*22*^ suggesting immature, rather than deficient, vasculature may be responsible for the higher hypoxia levels in Hs766t.^*2*^ At the cellular level, cell rupture and inflammation were evident in Hs766t but not in MiaPaCa-2 cells (Figs. 1B and D arrows). Despite their overall genetic similarity, the tumor microenvironment differs and Hs766t and MiaPaCa-2 can be easily distinguished by either anatomical MRI or histology.

**Figure 1:**
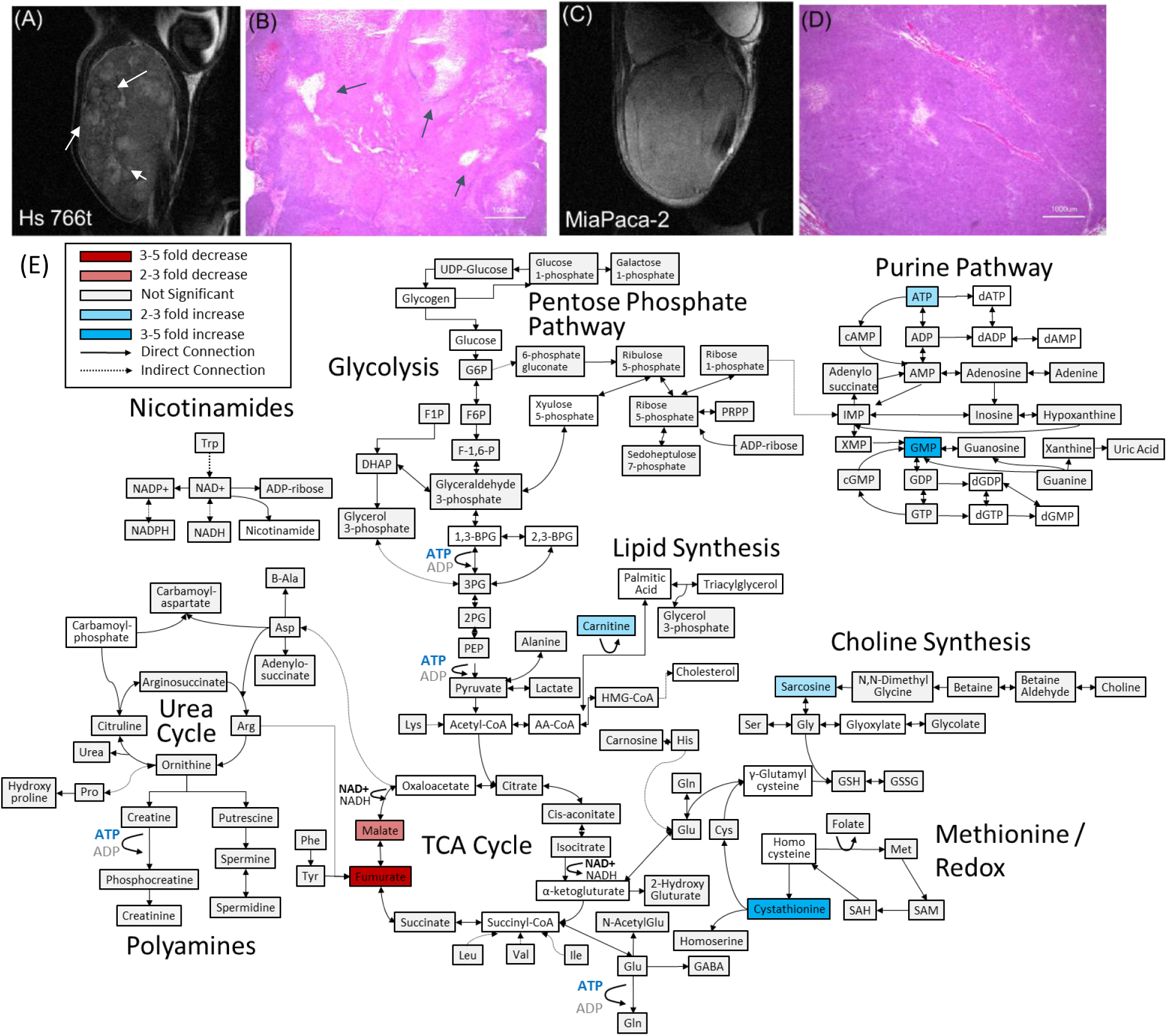
**(A and C)** T2 weighted anatomical RARE images of **(A)** HS766t and **(C)** PDAC xenografts implanted on the left leg. Focal necrosis is evident in the HS766t tumor, but not in the MiaPaCa-2 one. **(B and D)** H&E staining of biopsies from **(B)** HS766t and **(D)** MiPaca-2 tumors. Cell rupture is present in the Hs766t biopsy. **(E)** Metabolite differences of MiaPaCa-2 and Hs766t PDAC leg xenografts as analyzed by CE/MS. White boxes indicate metabolites not detected. Grey boxes indicate a statistically insignificant difference between cell lines (two-sided t-test, corrected for multiple comparisons by the two-stage linear step-up procedure of Holm et al with a confidence level of 5%^*66, 67*^). Blue and red boxes indicate statistically significant increases or decreases with respect to MiaPaCa-2.

### PDAC Xenografts Can Easily be Differentiated from Non-cancerous Host Tissue by Metabolic Differences

Both the MRI and histology results point to substantial differences in the tumor microenviroment between the two tumor types that may influence metabolism. Accordingly, we looked for alterations in central metabolic pathways for biosynthesis, stress response, and energetics that are commonly modified in tumor cell lines using capillary electrophoresis mass spectrometry (CE-MS) targeted metabolic profiling.^*23*^ CE-MS separates molecules by electrophoretic mobility. It has a different specificity than the more commonly used LC/MS, which separates molecules largely by size and charge rather than polarity. CE-MS displays greater sensitivity for small charged molecules such as amino acids compared to LC/MS, which is selective for large hydrophobic molecules such as lipids. CE/MS metabolic profiling can therefore give additional information not present in previous LC/MS experiments^*10*^.

As expected, pancreatic host tissue from the mouse was metabolically distinct from Hs766t and MiaPaca2 xenografts (p <0.001 based on two-way ANOVA), with numerous metabolic differences across multiple metabolic pathways (Fig. S2). The largest changes were concentrated in pathways connected to amino acid biosynthesis and degradation, reflecting an imbalance between amino acid metabolism and protein biosynthesis caused by unsustainable growth. Amino acid levels of all types and amino acid synthetic intermediates were strongly decreased in both cells lines. Both cell lines also show a strong depletion in the level of intermediates throughout the urea cycle, the primary pathway for protein catabolism, as well as the polyamine biosynthetic pathway downstream of the urea cycle.

Major metabolic changes are also evident in other pathways. Consistent with previous reports on PDAC tumors,^*24*^ flux through the glycolytic pathway is elevated in both cell lines but diverted into the pentose phosphate pathway for nucleotide and NADPH production. To counter the increased oxidative stress, there is increased activity through the methionine redox cycle: the reduced equivalents are depressed and the oxidized equivalents elevated compared to normal tissue. Finally, lactate levels are also highly elevated,^*25*^ consistent with a Warburg phenotype for both Hs766t and MiaPaCa-2.

### Fumarate Metabolism Distinguishes PDAC hypoxic subtypes

The metabolic differences between Hs766t and MiaPaca2 PDAC tumors were more subtle. Although it is possible to distinguish between the two types of PDAC tumors using the entirety of the metabolic profile (p=0.00015 for N=4, two-way ANOVA with Sidak’s correction for multiple comparisons), no single pathway stood out as being distinct (Fig. 1E). Only a few biomarkers are distinct at the 5% confidence level with most of the differences that do exist are in the TCA cycle. The most striking difference was in fumarate levels (p=0.003), which were significantly depressed (decreased 4 fold relative to normal) in MiaPaCa-2 and normal or slightly elevated in Hs766t (elevated 1.4 fold). Fumarate has been suggested as an oncometabolite^*26, 27*^ created both through the TCA cycle and as a byproduct of the urea cycle that competitively inhibits 2-OG dependent oxygenases to stabilize the HIF complex and induce pseudohypoxia. Malate and arginosuccinate, two other intermediates in the fumarate pathway were also significantly depressed in MiaPaCa-2.

### Pyruvate metabolism is indistinguishable between PDAC hypoxic subtypes

The CE/MS experiment measures the static distribution of metabolites within the tumor, which is the sum of multiple biochemical pathways. While the data suggests that a difference in glycolysis and oxidative phosphorylation may exist between the MiaPaCa-2 and Hs766 cell lines, the statistical significance of these changes is mostly uncertain and the origin of the effect is not clear - it is uncertain whether the difference is the result of upregulation of specific genes or is a more general effect from changes in the underlying physiology of the tumor microenvironment. To more directly probe specific enzyme activities within the glycolytic and TCA cycles, we first tracked the *in vivo* utilization of hyperpolarized ^13^C labelled pyruvate using magnetic resonance spectroscopy to detect the *de novo* generation of new metabolites from pyruvate. Pyruvate metabolism is a central control point between glycolysis and oxidative phosphorylation and dysregulation of pyruvate dehydrogenase can be an important component of the Warburg effect.^*28*^ Quantifying pyruvate to lactate exchange through a ^13^C tracer provides an estimate of flux through the TCA, Cahill, and anaerobic fermentation pathways, and can provide a measurement of the degree of dysregulation of central metabolism.^*29*^

Figs. 2A and B shows typical spectra after the injection of 98 mM solution of hyperpolarized [1-^13^C]pyruvate into the tail vein of nude mice bearing MiaPaCa-2 or Hs766t xenografts in the left leg. The five peaks correspond to pyruvate (172.6 ppm), lactate (184.9), alanine (178.2), bicarbonate (162.6 ppm), and inactive pyruvate hydrate (180.9 ppm). Few differences could be seen when using C-1 labeled pyruvate as a metabolic tracer; pyruvate metabolism appears to be statistically indistinguishable in the MiaPaCa-2 and Hs766t cell lines. The rate of pyruvate to lactate conversion was similar in MiaPaCa-2 and Hs766t as was the flux through the Cahill cycle to alanine and the first step of the TCA cycle as measured by bicarbonate production (see Fig. 2). While differences in pyruvate metabolism in hypoxic and oxidative SU8686 tumors has previously been shown by hyperpolarized C-1 labeled pyruvate, pyruvate metabolism is not a sensitive biomarker for distinguishing subtle differences among hypoxic pancreatic adenocarcinoma subtypes.

**Figure 2:**
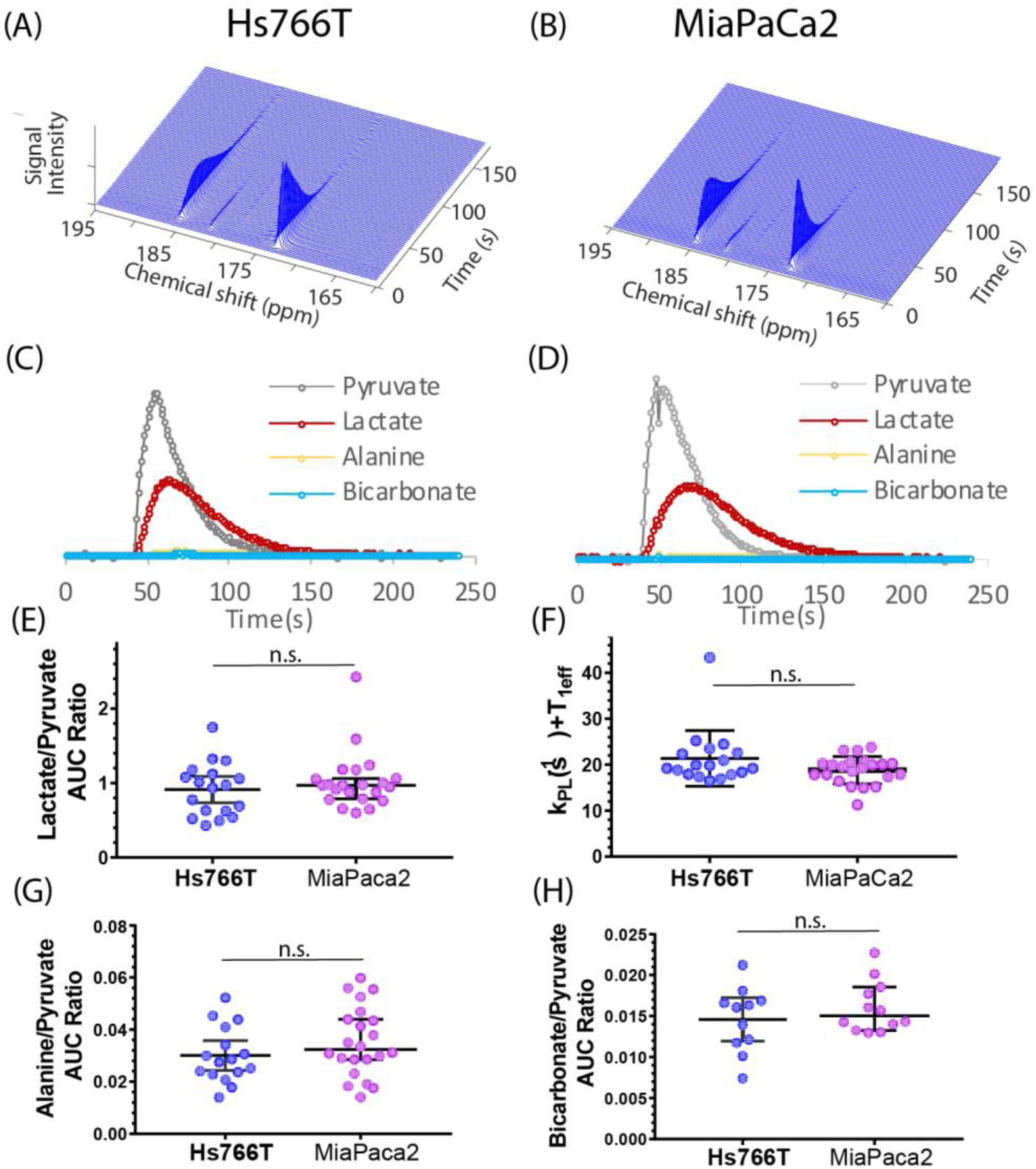
Pyruvate metabolism is similar in Hs766T and MiaPaca2 Xenografts. **(A and B)** Representative signal after injecting 300 µL of 98 mM hyperpolarized [1-^13^C]pyruvate into the tail vein of a mouse of a nude mouse with either a **(A)** Hs766T or **(B)** MiaPaCa2 leg xenograft. Signal loss is due to a combination of the loss of hyperpolarization and conversion of pyruvate to other metabolites. Corresponding kinetic traces of the pyruvate, lactate, alanine and bicarbonate signals metabolites for **(C)** Hs766T and **(D)** MiaPaCa2 xenografts. **(E)** Ratio of the integrated lactate and pyruvate for Hs766T (n=18) and MiaPaca2 (n=22) mice. The ratio is equal to the net lactate to pyruvate conversion rate in the absence of lactate efflux or back conversion. **(F)** Decay rate of the pyruvate signal, equivalent to the sum of the net lactate to pyruvate conversion rate and the effective relaxation rate (T1eff, assumed to be the same between cell lines). **(G and H)** Ratio of the integrated alanine (**G**) or bicarbonate **(H)** to pyruvate. No statistically significant difference between cell lines was detected for any measure (Mann-Whitney rank test). Error bars represent 95% confidence intervals.

### Glucose metabolism, but not glucose uptake, distinguishes PDAC hypoxic subtypes

The failure of hyperpolarized [1-13C] pyruvate encouraged us to look elsewhere for possible metabolic biomarkers. The CE/MS data is suggestive of an upregulation in MiaPaCa-2 of the later stages of glycolysis relative to Hs766t, but the sample-sample variability inherent to MS techniques obscures the magnitude of any difference (see Fig. S3). Hyperpolarized ^13^C MRS is more precise in this respect, but the transient nature of hyperpolarization restricts analysis to only the first few metabolic steps away from the probe.^*17*^ To probe the glycolytic pathway, a different technique is needed.

We have previously shown that it is possible to use the correlation of the ^13^C signal in both time and space to reduce the noise level in the signal by an order of magnitude or more (see Methods).^*17*^ Using this technique, we first checked the glucose metabolism of each tumor type following an injection of 50 mg bolus of [U-^13^C]glucose using non-localized spectroscopy. The resulting spectra are complex and include contributions from the α and β forms of glucose, the endogenous lipid signal, and the signals from the end products of anaerobic fermentation and the Cori and Cahill cycle lactate and alanine (Fig. 3A). The peak at 95 ppm is of particular interest as it can be assigned to specifically glucose and glucose-6-phosphate without contributions from other glycolytic intermediates. Following the intensity of the peak at 95 ppm therefore gives an indication of the first steps of glycolysis, glucose import and phosphorylation. The other major glucose peak at 60 ppm contains contributions from all glycolytic intermediates and serves as a measure of the overall progress of glycolysis. Lactate and alanine are observed as partially resolved shoulders of the broad lipid peak at 19.8 ppm and 16.8 ppm, respectively.

**Figure 3:**
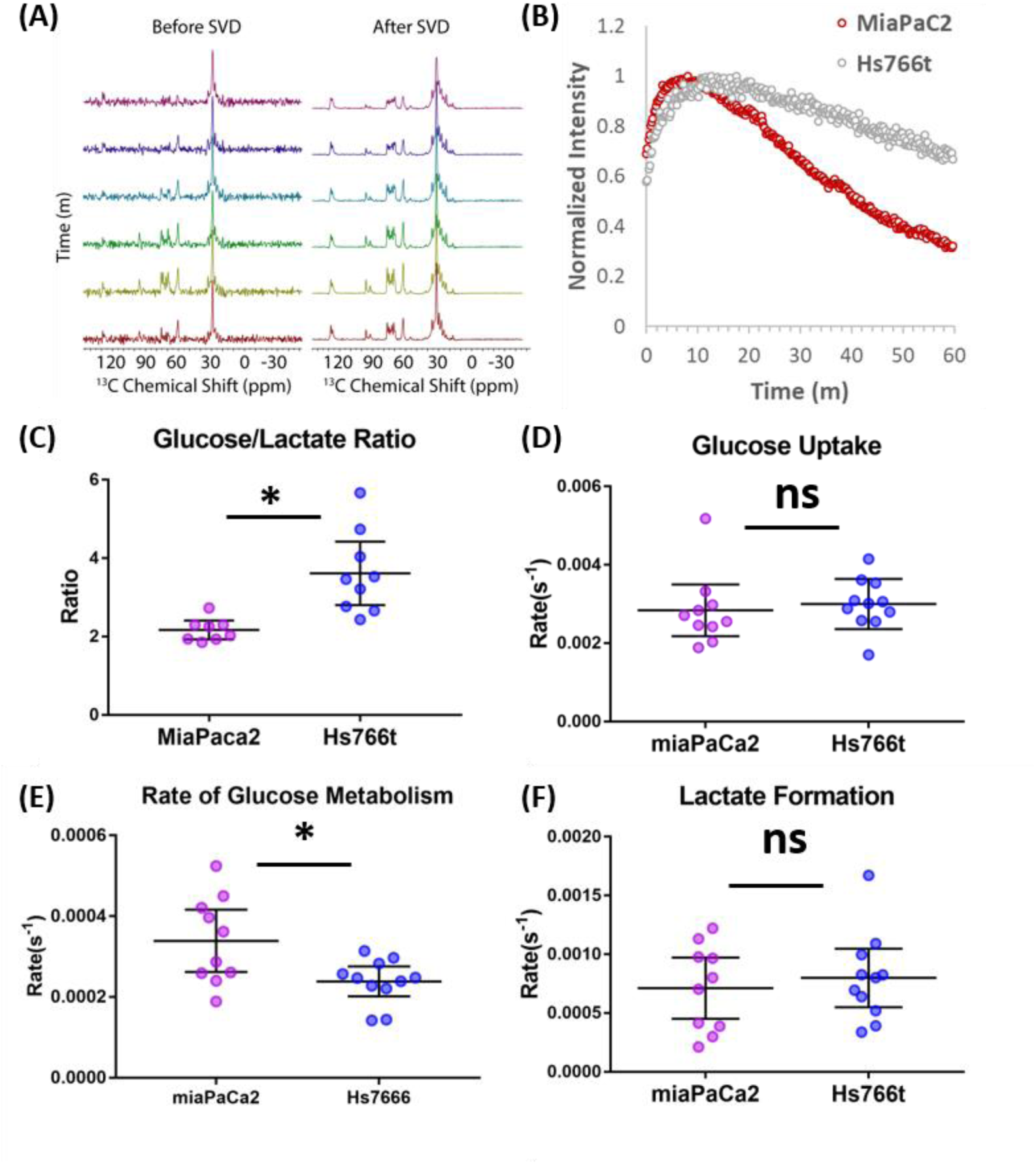
Glucose Metabolism Differentiates Hs766T and MiaPaca2 Xenografts. **(A)** Representative signal before and after rank reduction by SVD after injecting 300 µL of 98 mM hyperpolarized [1-^13^C] glucose into the tail vein of a mouse of a nude mouse with a MiaPaCa-2 leg xenograft. The largest signal at 29 ppm is due to endogenous lipids, while glucose peaks can be seen in region from 60-100 ppm and lactate and alanine peaks can be detected at 23 ppm and 19 ppm, respectively. No hyperpolarization is used in this experiment; the time dependence of the signal is due to metabolic interconversion. **(B)** Representative kinetic traces for the glucose signal for Hs766T (n=10) and MiaPaCa-2 (n=11) xenografts. **(C)** Whether expressed as either a ratio or directly as rate obtained from curve-fitting (E) a statistically significant difference in the rate of glucose metabolism can be seen between the Hs766t or MiaPaCa-2 PANC subtypes (Mann Whitney rank test, p=0.03 and 0.02 respectively). No statistically significant difference could be seen in the rate of glucose uptake **(D)** or lactate production **(F).** Error bars represent 95% confidence intervals.

The overall kinetics of glycolysis from following the 60 ppm peak approximately matched previous ^13^C measurements of glycolysis (Fig. 3B). However, the improvement in temporal resolution (5 m vs 12 s) afforded by greatly increased signal to noise allows an assessment of the fast glucose import step by MRI, which could not be resolved effectively before. No difference between cell lines could be detected in the rate of glucose uptake (Fig.3D), in agreement with the similar levels of the glucose transporter GLUT1, detected by western blot (see Fig. S1), or in the rate of lactate formation (Fig. 3F). The rate of glucose metabolism after import; on the other hand, distinguishes MiaPaCa-2 and Hs766t xenografts. Hs766t xenografts displayed a statistically significant slower glucose metabolism than MiaPaCa-2 xenografts (Fig. 3E, Mann-Whitney rank test, p=0.02). This difference is also reflected in the time-averaged glucose to lactate ratio (Fig. 3C, Mann-Whitney rank test, p=0.04); however, interpretation of the ratio as a rate^*30*^ is complicated by the contribution of the endogenous lipid signal to peak.

### Local differences in metabolism exist within PDAC tumors

Figure 1E shows that a metabolomic profile from a CE/MS can distinguish between xenografts of the MiaPaCa-2 and Hs766t cell lines. Unfortunately, mass spectrometry can only be done *ex situ* and it is impossible to apply this procedure to see the distribution of metabolites in a living tumor. MRI spectroscopy can see the distribution but cannot distinguish the source. The higher signal to noise afforded by the rank reduction technique allowed the direct determination of the kinetics of glucose metabolism *in vivo* by MRI. A multidimensional extension of the technique, tensor decomposition, may open up the possibility of metabolic imaging. As a first test, we used chemical shift imaging to provide a low-resolution (1.5×1.5×16 mm) map of the rates of glycolysis and anaerobic fermentation (Fig.4 A-D). Simple chemical shift imaging was used to minimize potential imaging artifacts; considerable acceleration can be achieved by a more efficient pulse sequence and is the focus of ongoing research.

Figure 4 shows representative results from chemical shift imaging of MiaPaCa-2 and Hs766t xenografts before and after noise suppression (see Methods). While the raw images are mainly noise, (Fig.4 E) the processed images by tensor decomposition clearly show localized uptake of glucose within the tumor. As in the non-localized experiment, the glucose signal can be seen to decay and the corresponding lactate signal at 19 ppm to simultaneously increase as the tumor metabolized the bolus (Figure 4G). Local differences in metabolism can be detected in many tumors. For example, in one Hs766t xenograft (Figure 4G) glucose metabolism is distributed relatively uniformly after taking into account the overall tumor anatomy. Lactate production, on the other hand, is localized in this tumor to one side where focal necrosis is more evident. In comparable MiaPaCa-2 tumors (Fig. 5), glucose and lactate production appears to be more tightly correlated, congruent with the greater homogeneity apparent in the anatomical MRIs.

**Figure 4:**
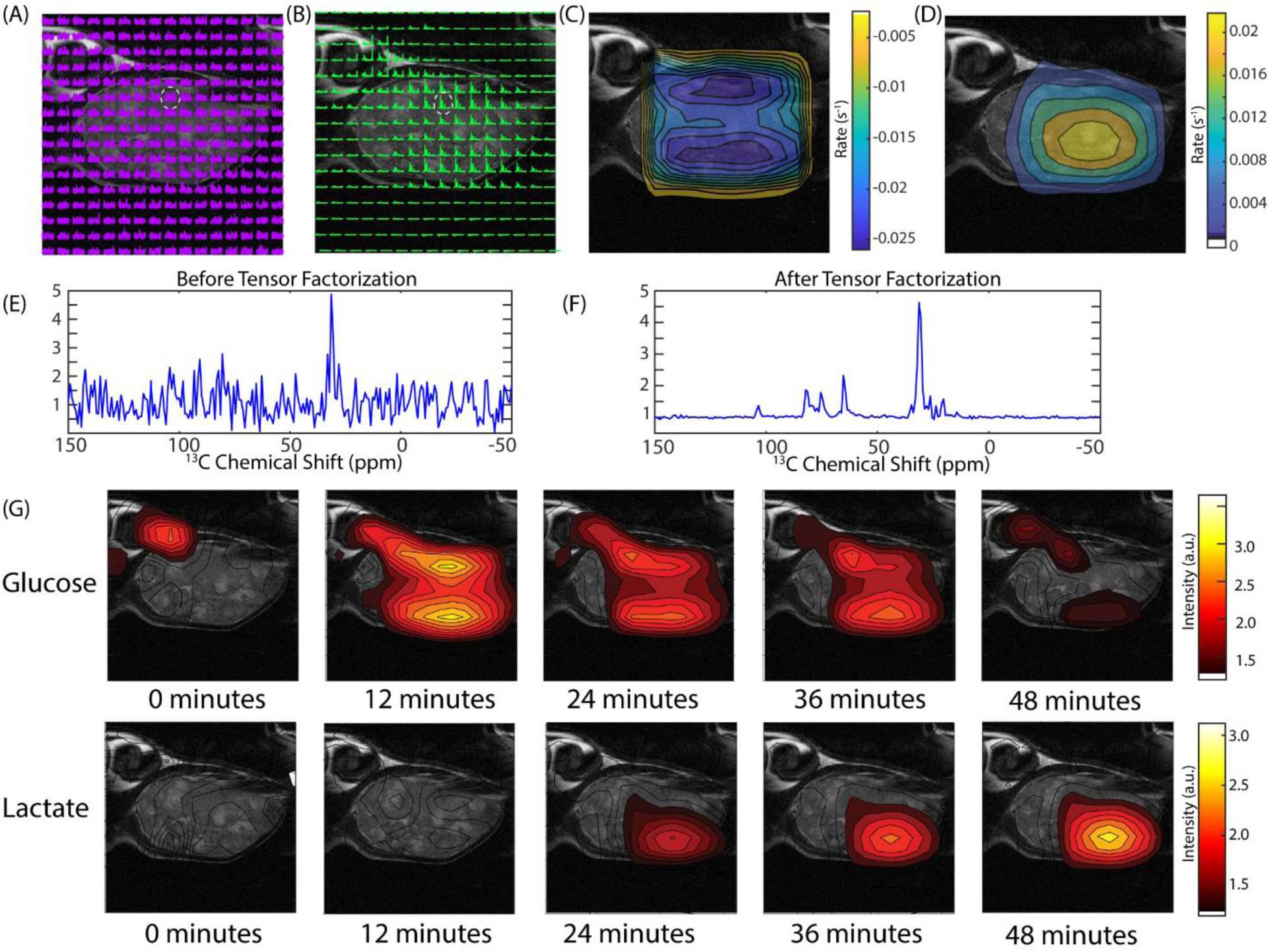
CSI imaging of a Hs766t mouse leg xenograft after a 50 mg [U-^13^C] glucose injection in a volume of – microliters of PBS. An 8×8 image of the tumor bearing mouse leg was acquired by chemical shift imaging every 48 s for 60 min. The final image was zero-filled to 16×16. Each is voxel 0.15 cm × 0.15 cm × 1.6 cm in size. **(A)** The glucose region of the spectra at 12 min overlaid on the anatomical image. **(B)** Same image after tensor factorization. **(C and D)** Rate map of **(C)** glucose and **(D)** lactate metabolism calculated from the image series **(E and F).** Signal from the voxel indicated by the white dashed line **(E)** before and **(F)** after tensor factorization. **(G)** Contour maps created from the peak maximums of the glucose and lactate signals at the time points indicated.

**Figure 5:**
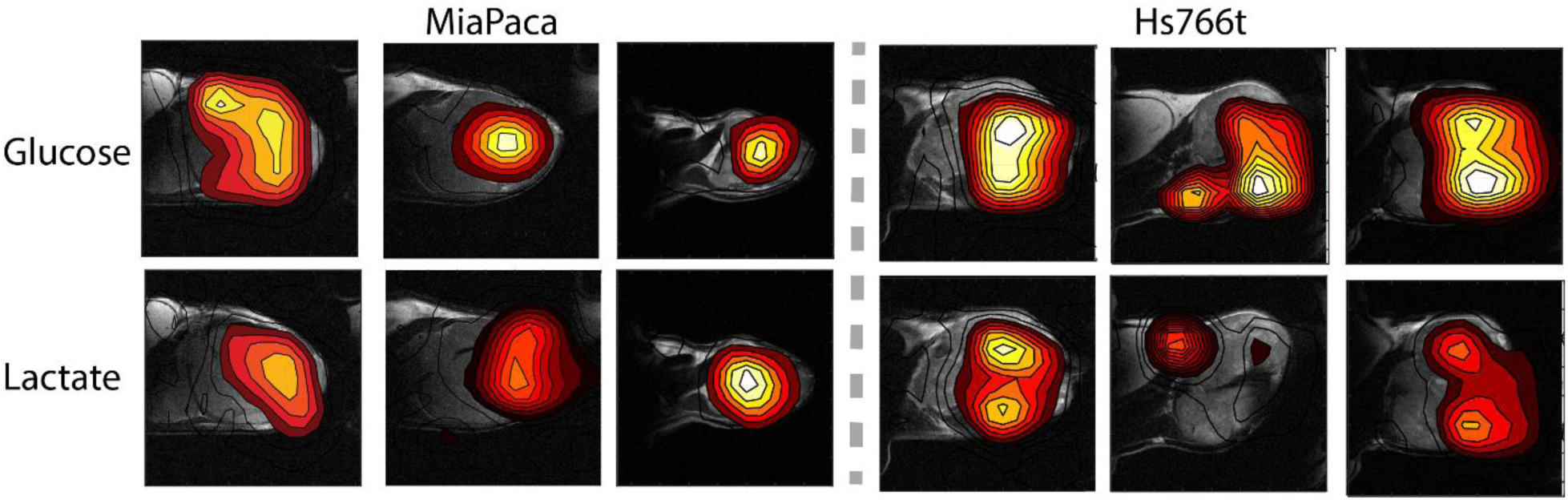
Contour maps created from time averages of the peak maximums of the glucose and lactate signals for three representative MiaPaCa-2 (left) and Hs766t (right) tumors.

## Discussion

Pancreatic ductal adenocarcinoma (PDAC) represent 90% of pancreatic cancers and are characterized by a poor prognosis and limited treatment strategies.^*31, 32*^ Given PDACs resistance to traditional chemo-and radiotherapy regimes,^*33*^ alternative points of attack are being considered. One potential point of attack is the dysregulated metabolism of PDACs,^*34*^ which is highly dependent on protein autophagy and catabolism^*35, 36*^ and exogenous glutamine and glucose.^*24, 37*^ Further, PDAC tumors usually have alterations in the activity in the urea cycle to support pyrmidine and amino acid synthesis^*38, 39*^ and often display a Warburg phenotype of increased glycolysis followed by diversion to lactate.^*40*^ Each alteration and dependency represents a potential point of intervention. Although targeting the master genetic switches for these transformations, p53 and kRAS,^*24, 41*^ is difficult, the downstream enzymes are practical targets. Inhibitors for lactate dehydrogenase^*42*^ and the lactate transporter MCT1^*43*^ have shown promise in preclinical trials and may enter clinical trials in the near future. Beyond the Warburg effect, researchers have begun to target other vulnerable aspects of PDAC metabolism such as amino acid synthesis^*44*^ or the unique chemical environment of tumors by hypoxia activated prodrugs.^*2, 45*^

Targeting aberrant metabolism requires a method of monitoring treatment progress and selecting suitable patient populations. Response to cancer treatment can be highly variable and there is a concerted push to tailor treatment regimes to individual patients.^*46*^ For protein targets such as receptors, genome sequencing or protein expression profiling is often sufficient to demonstrate a patient has a vulnerable mutation. Targeting aberrant metabolism is more difficult as the metabolism of tumors is not limited to the tumor itself, but contains substantial contributions from the surrounding cells both directly through diffusion of metabolites across the tumor boundary^*47*^ and indirectly through the influence of regulatory and epigenetic signals.^*48*^ The physical microenvironment of the tumor can also affect metabolism. Deficient or improperly formed^*2*^ vasculature often induces hypoxia in PDAC tumors^*49*^ which can induce metabolic changes^*23, 50*^ that would not be evident by genetic analysis alone.

The steady state metabolism of PDAC tumors can be probed indirectly through analysis of urine, blood, or pancreatic cyst fluid^*51*^ or more directly through magnetic resonance spectroscopy. However, metabolic networks are more flexible than protein networks and flux through the network can be rerouted to limit the impact of targeted enzymes.^*37*^ Evaluating the target engagement of potential inhibitors can be difficult under these conditions. We demonstrate the potential of multimodal metabolic profiling of PDAC tumors for distinguishing animal models that are genetically similar but display very different phenotypes. Hs766t is a cell line derived from lymphatic metastasis of pancreatic cancer that generates highly necrotic, hypoxic, slow growing heterogenous tumors. MiaPaCa-2 is derived from primary cancer whose tumors are less necrotic, grow faster, and are highly homogenous. Despite their dissimilar origin and physiological differences, the steady state metabolism probed by CE/MS of Hs766t and MiaPaCa-2 is fairly similar with only a few potential differentiating biomarkers (Fig. 1E). Hyperpolarized pyruvate-lactate fluxes of Hs766t and MiaPaCa-2 xenografts estimated by ^13^C hyperpolarized pyruvate MRI were statistically indistinguishable (Fig. 3). The flux through the TCA cycle and anaerobic respiration is similar, as expected from the presence of KRAS and P53 mutations in both cell lines and similar LDHA levels (Fig. S1).

Using [U-^13^C] glucose instead of [1-^13^C] pyruvate allowed a more encompassing overview of metabolism. Measurements of in vivo glucose metabolism by ^13^C MRI have proved difficult because of difficulty of hyperpolarizing glucose and the low SNR in non-hyperpolarized experiments. The recently developed a post-processing novel denoising algorithm recovered sufficient SNR from non-hyperpolarized MRI imaging experiments of glucose metabolism.^*17*^ The most important information acquired by this method is the direct measurement of rates of glycolysis and anaerobic respiration, allowing imaging of the Warburg effect. High glucose uptake is one of the most well studied features of cancer and has been utilized in FDG-PET imaging in clinical settings. FDG-PET is limited in that the radiotracer cannot differentiate among the later steps of glycolysis. By allowing detection further down metabolic pathways than FDG-PET, investigating glucose metabolism by ^13^C MRI can potentially probe more subtle defects. We see a similar effect in Fig 3; among Hs766t and MiaPaCa-2 PDAC xenograft animal models, glucose uptake is similar but glucose metabolism is distinct.

These results suggest some advantages for ^13^C glucose imaging. Compared to FDG-PET, there is no need for a radioactive tracer, which makes this imaging potentially safer and less invasive. By observing the lactate production at later time points, ^13^C glucose imaging can potentially detect cancer even in highly glucose consuming tissue such as brain or liver. It can also potentially detect cancer in the bladder because glucose is not excreted immediately in urine, while FDG excreted in the urinary tract and excess signal in the bladder can interfere with lesion detection within or near the bladder wall. In comparison to hyperpolarized MRI, non-hyperpolarized ^13^C glucose imaging does not require onsite preparation of the probe, removing one of the main barriers to clinical translation of metabolic imaging by MRI. While some challenges remain and the technique is inferior to PET in some respects, particularly with respect to resolution and imaging time, ^13^C glucose imaging by MRI may emerge as a viable adjunct or alternative to FDG-PET.

## Materials and Methods

### Mouse Models

The animal experiments were conducted according to a protocol approved by the Animal Research Advisory Committee of the NIH (RBB-159-2SA) in accordance with the National Institutes of Health Guidelines for Animal Research. Female athymic nude mice weighing approximately 26 g were supplied by the Frederick Cancer Research Center, Animal Production (Frederick, MD) and housed with *ad libitum* access to NIH Rodent Diet #31 Open Formula (Envigo) and water on a 12-hour light/dark cycle. Xenografts were generated by the subcutaneous injection of 3 ×10^6^ MIA PaCa-2 (America Type Cell Collection (ATCC), Manassas, VA, USA) or Hs766t (Threshold Pharmaceuticals, Redwood City, CA, USA) pancreatic ductal adenocarcinoma cells.^*52*^ Both cell lines were tested in May 2013 and authenticated by IDEXX RADIL (Columbia, MO) using a panel of microsatellite markers.

### CE/MS analysis

Tumors were excised when the volume reached 600 mm^3^ and immediately frozen in liquid nitrogen and stored at −80 °C until analysis. A total of 116 metabolites involved in glycolysis, the pentose phosphate pathway, the tricarboxylic acid cycle, the urea cycle, and polyamine, creatine, purine, glutathione, nicotinamide, choline, and amino acid metabolism were analyzed using CE-TOF and QqQ mass spectrometry (Carcinoscope Package, Human Metabolome Technologies, Inc.)

### Western Blotting Analysis

The mice bearing MiaPaCa-2 and Hs766t tumors (n = 4 for each group) were euthanized by breathing carbon dioxide gas, and tumor biopsy samples were excised. The tumor tissues were immediately homogenized with T-PER tissue protein extraction reagent (Thermo scientific). The homogenate was centrifuged, and the supernatant was used for Western blot analysis. Hexokinase-2, Glut-1, LDHA proteins in tumor extract were separated on 4% to 20% Tris-Glycine gel and CD31 was separated on NuPAGE 3 to 8 % Tris-Acetate gel (Life Technologies) by SDS-PAGE and were transferred to nitrocellulose membrane. The membranes were blocked for 1hour in blocking buffer (3% nonfat dry milk in 0.1% Tween 20/TBS), which was then replaced by the primary antibody (1:500-1:1,000) diluted in blocking buffer, and were incubated for 1 hour at room temperature. The membranes were then washed three times in washing buffer (0.1% Tween 20/TBS). The primary antibody was detected using horseradish peroxidase–linked goat anti-mouse or goat anti-rabbit IgG antibody at a 1:2,000 dilution (Santa Cruz Biotechnology), visualized with Western Lightning Plus-ECL enhanced chemiluminescence substrate (Perkin Elmer Inc.) and measured by the Fluor Chem HD2 chemiluminescent imaging system (Alpha Innotech Corp.). Density values for each protein were normalized to actin or HSC70.

### ^13^C MRS with hyperpolarization

Samples for NMR were prepared and analyzed as previously described in Ref. 17. [1-^13^C]pyruvic acid (30 μL), containing 15 mM TAM and 2.5 mM gadolinium chelate ProHance (Bracco Diagnostics, Milano, Italy), was hyperpolarized at 3.35 T and 1.4 K using the Hypersense DNP polarizer (Oxford Instruments, Abingdon, UK) according to the manufacturer’s instructions. Typical polarization efficiencies were around 20%. After 40-60 min, the hyperpolarized sample was rapidly dissolved in 4.5 mL of a superheated HEPES based alkaline buffer. The dissolution buffer was neutralized with NaOH to pH 7.4. The hyperpolarized [1-^13^C]pyruvate solution (96 mM) was intravenously injected through a catheter placed in the tail vein of the mouse (1.1 mmol/kg body weight). Hyperpolarized ^13^C MRI studies were performed on a 3 T scanner (MR Solutions, Guildford, UK) using a home-built ^13^C solenoid leg coil. After the rapid injection of hyperpolarized [1-^13^C] pyruvate, spectra were acquired every second for 240 s using a single pulse acquire sequence with a sweep width of 3.3 kHz and 256 FID points.

### Dynamic ^13^C Glucose MRS without hyperpolarization

The magnetic resonance spectroscopy experiments were performed on either a 9.4 T Biospec 94/30 horizontal scanner or a MR Solutions 3 T horizontal scanner using a 16 mm double resonance 1H/13C coil constructed as described in Ref. 17. Each mouse was anesthetized during imaging with isoflurane 1.5–2.0% administered as a gaseous mixture of 70% N2 and 30% O2 and kept warm using a circulating hot water bath. Both respiration and temperature were monitored continuously through the experiment and the degree of anesthesia adjusted to keep respiration and body temperature within a normal physiological range of 35-37° C and 60-90 breaths per min. Anatomical images were acquired with a RARE fast spin echo sequence^*53*^ with 15 256×256 slices of 24 mm × 24 mm × 1 mm size with 8 echoes per acquisition, a 3 s repetition time, and an effective sweep width of 50,000 Hz. Samples were shimmed to 20 Hz on the 9.4 T with first and second order shims using the FASTMAP procedure.^*54*^ Non-localized spectra of [U-1^3^C] glucose without DNP at 9.4 T were acquired with the NSPECT pulse-acquire sequence using maximum receiver gain, a repetition time of 50 ms, Ernst Angle excitation of 12°, 256 FID points, a sweep width of 198.6 ppm, 16 averages per scan, and 4500 scans for a total acquisition time of 1 hour. MLEV16 decoupling^*55, 56*^ was applied during acquisition using −20 dB of decoupling power and a 0.2 ms decoupling element. The decoupling pulse was centered on the main proton lipid resonance at 1.3 ppm. Chemical shift imaging experiments were performed similarly except an 8×8 image using 0.3 cm × 0.3 cm × 1.5 cm voxels was acquired every 48 seconds (4 averages per scan) for 90 minutes.

### Signal processing

For non-localized (two dimensional) experiments, the first 67 points of the FID in the time dimension were removed to eliminate the distortion from the group delay corresponding to the 13 ms dead time of the Bruker 9.4 T.^*57*^ The FID was Fourier transformed and the phase estimated by the entropy minimization method of Chen et al,^*58*^ as implemented in MatNMR.^*59*^ After low rank reconstruction by SVD (see below), the baseline was estimated by a modification of the Dietrich first derivative method to generate a binary mask of baseline points,^*60*^ followed by spline interpolation using the Whittaker smoother^*61*^ to generate a smooth baseline curve.^*62*^ The final correction adjusts for the limited number of points in the frequency dimension by continuation of the FID by linear prediction. The remaining 189 points of the FID after truncation in the first step were extrapolated to 1024 points using the “forward-backward” linear prediction method of Zhu and Bax.^*63*^ Fourier transforming the FID of the transients from each voxel individually generated the final spectrum. Phase estimation proved difficult for to the chemical shift imaging experiments and therefore the spectra for chemical shift imaging experiments are shown in magnitude mode.

### Low Rank Reconstruction

For the two-dimensional signal matrices generated by non-localized pulse acquire experiments, the rank reduced signal was generated by truncating the SVD by setting the *N-r* diagonal values of the singular value matrix *S* to 0, where *N* is the number of rows in *S* and *r* is the predicted rank. The predicted rank was set to 5 unless otherwise specified, which is equal to the number of independent species in the hyperpolarized pyruvate experiment. Tensor decomposition was achieved through higher order orthogonal iteration^*64*^ in the Matlab NWay package^*65*^ using a rank of 8 in the temporal and spatial dimensions and 6 in each spatial dimension.

## Supplementary Figures

**Figure S1:**
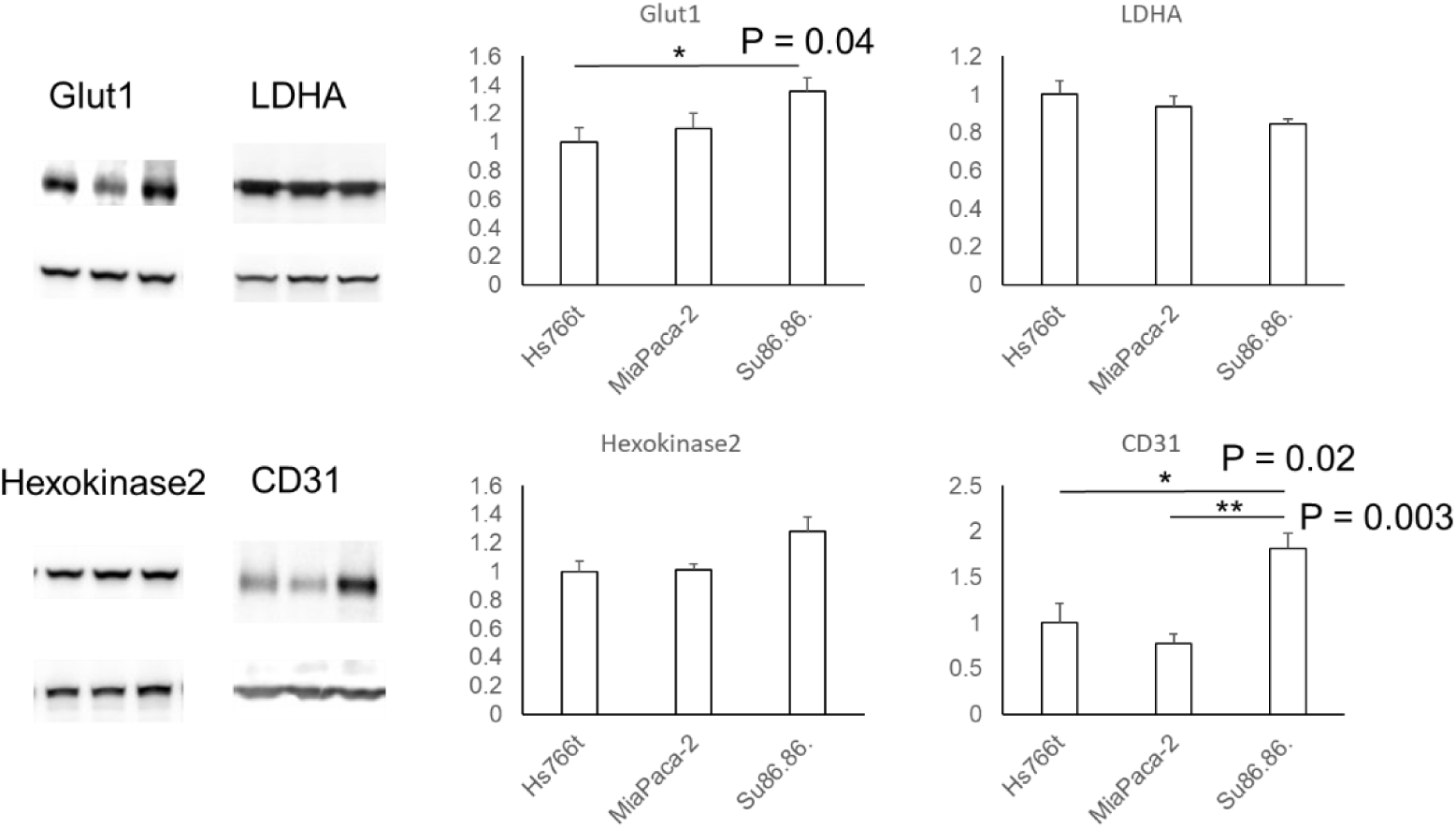
Protein expression levels from immunoblotting of tumor extracts of key proteins associated with metabolism. Su86.86 forms a distinct subtype with statistically significant differences of the glucose transporter 1 and the angiogenic factor CD31. Error bars represent standard deviation (n=4).

**Figure S2:**
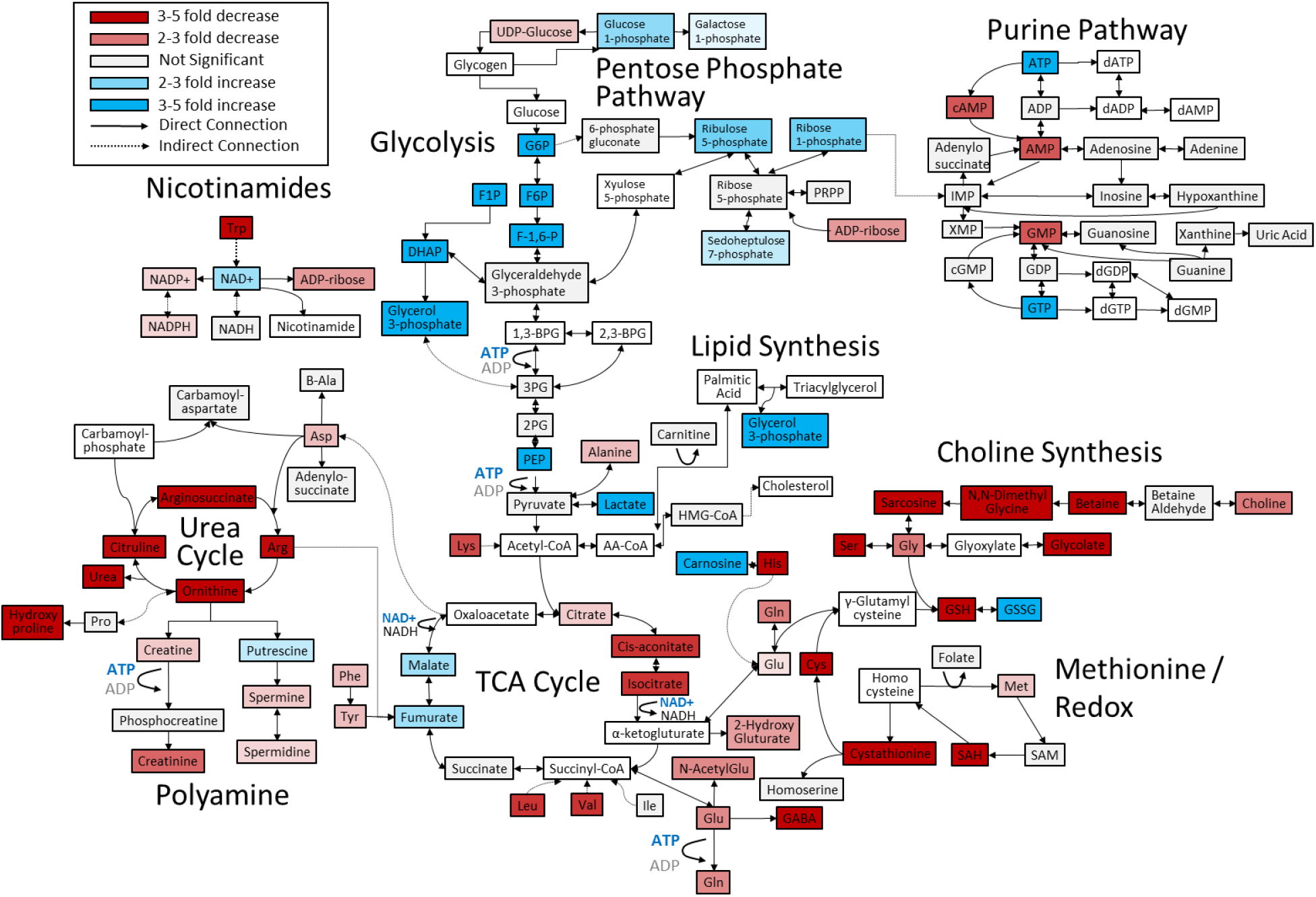
Metabolite differences from normal tissue compared to the MiaPaCa-2 PDAC leg xenografts as analyzed by CE/MS. White boxes indicate metabolites not detected. Grey boxes indicate a statistically insignificant difference between cell lines (two-sided t-test, corrected for multiple comparisons by the two-stage linear step-up procedure of Holm et al with a confidence level of 5%^*66, 67*^). Blue and red boxes indicate statistically significant increases or decreases with respect to normal tissue.

**Figure S3:**
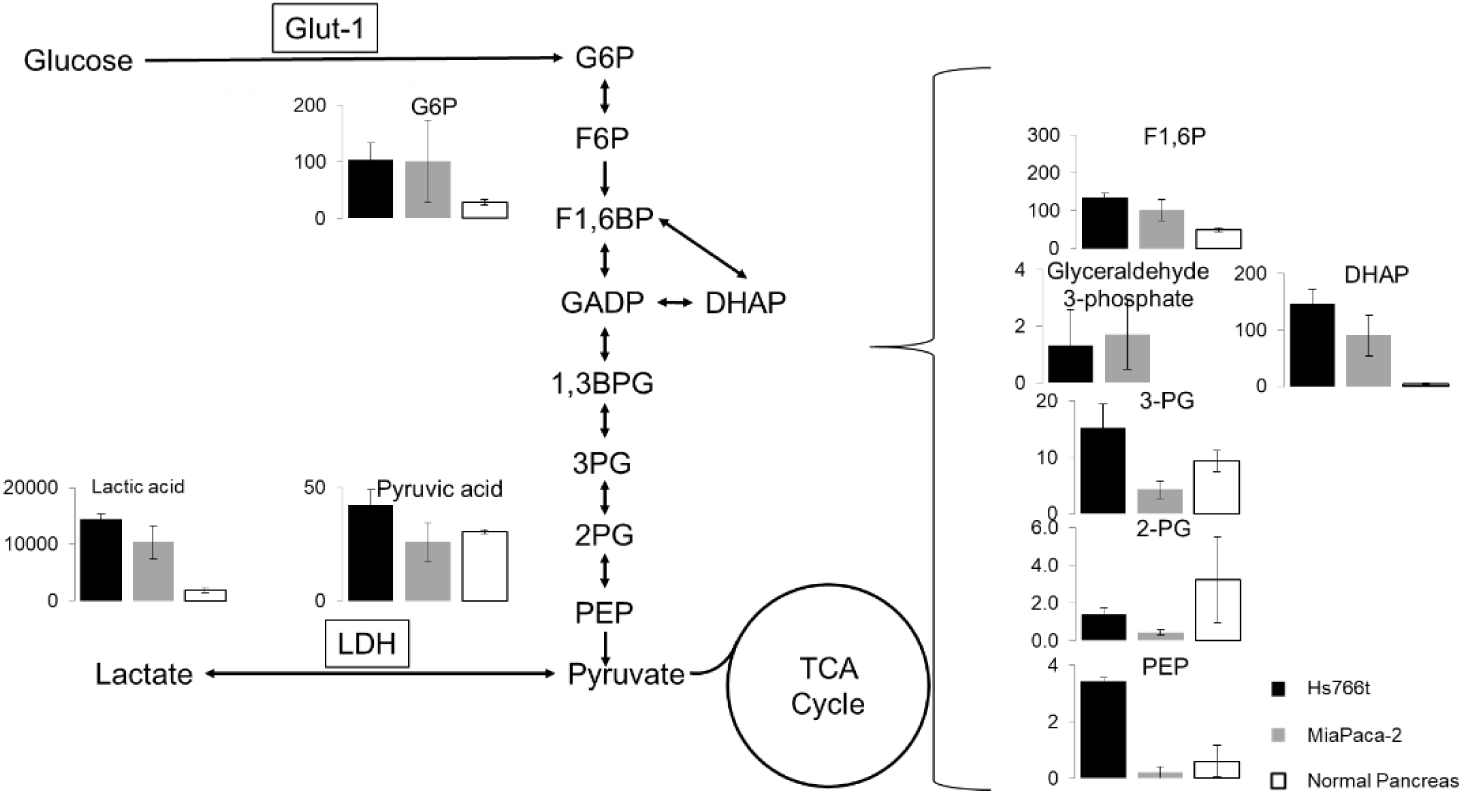
Metabolic differences within glycolysis between normal tissue and MiaPaCa-2 and Hs766t PDAC leg xenografts. Differences between normal tissue and cancerous can be seen throughout. Hs766t and MiaPaCa-2 diverge after the pentose phosphate shunt at glyceraldehyde-3-phosphate.

